# Risk Factors and Patterns of *Treponema* Infection Affecting Olive Baboons in Gombe National Park

**DOI:** 10.1101/2025.11.07.687138

**Authors:** Dismas Mwacha, Anthony Collins, Jane Raphael, Philemon Wambura, Abubakar Hoza

## Abstract

*Treponema pallidum* subsp. *pertenue* (*TPE*) causes yaws, a chronic, nonvenereal treponematosis characterized by contagious cutaneous lesions in early stages and destructive bone involvement in the tertiary stage. In the latency stage, the infection is asymptomatic with only serologic markers.

A retrospective analysis of health, demographic, and behavioral records using a Generalized Linear Mixed Model (GLMM) with binomial distribution, conducted through R software to assess disease risks and trends from January 2019 to December 2024 of wild baboons at Gombe, revealed an overall clinical prevalence of 11.24% *(2018/17946*) across the eight studied troops. Age was a strong predictor where infants (OR = 0.05, p < 0.001), juveniles (OR = 0.27, p < 0.001), and subadults (OR = 0.65, p < 0.01) had significantly reduced odds of displaying *Treponema* signs compared to adults. Troops B.C and D.D exhibited elevated infection risk (OR = 2.74 and 2.63, respectively). Pregnant females (OR = 0.19, p < 0.001), wounded baboons (OR = 0.45, p < 0.001), and male immigrants (OR = 0.47, p < 0.001) were less likely to show signs. Infection signs were also lower during wet seasons (OR = 0.81, p < 0.001). Notably, the odds of infection increased consistently over time (Year OR = 1.73, p < 0.001).

Understanding the ecological and demographic determinants of *Treponema* transmission is essential for disease surveillance, conservation, and One Health initiatives. This study presents the first long-term dataset on (TPE) infection in wild baboons from Tanzania and East Africa.

**Author Summary:** *Treponema pallidum subspecies pertenue (TPE)* bacteria can cause debilitating disease in both humans and animals, yet little is known about how it spreads among wild Non-Human Primates. We studied a large population of habituated baboons and analyzed how various factors like Age, Troop, Pregnancy, Wounds, male immigration, Treatments and Interactions affect the likelihood of catching and showing signs of *Treponema* infection. Our analysis found that younger baboons are much less likely to show symptoms, but infection risk increases with Age. Some troops had significantly higher infection rates compared to others. Pregnant females and immigrants (Males) were less likely to show symptoms, possibly due to behavioral or biological differences. The risk of infection has also been increasing year after year. These findings suggest *Treponema* is becoming more common and highlight the importance of targeted monitoring and control in high-risk groups. Understanding these dynamics helps us protect wild primates and prevent cross-species transmission and potentially spill over to humans.

## Introduction

Gombe National Park is famous for chimpanzees *(Pan troglodytes schweinfurthii)*, Jane Goodall began her studies in the area in 1960, and in 1964, the Gombe Stream Research Centre (GSRC) was established. Gombe’s research focuses on life histories, tracing individuals, their families, and their communities over time.

Baboons *(Papio anubis)* have been studied at Gombe since 1967, in parallel to the chimpanzee study but recording less detail on a much larger sample of habituated baboons. Starting with 93 baboons in two groups in 1967, groups grew and divided until reaching eleven groups by 1999 (325 recognized baboons), of which currently has 8 troops totaling about 209 baboons are under study. Most of the recorded baboon data comprises births, deaths, male intergroup migrations, illnesses, injuries, monthly diet, interactions with other species, interactions between different groups, and any unusual events [1–3].

*Treponema* diseases are caused by a group of spiral-shaped bacteria known as *Treponema*, which include several subspecies that affect humans and animals. The most well-known disease caused by these bacteria is syphilis (*Treponema*, subsp. *pallidum* (TPA), which is primarily transmitted through sexual contact. Syphilis is characterized by four (4) stages, namely primary stage (characterized by a painless sore), secondary stage (characterized by skin rashes, fever, etc.), latent stage (no specific symptoms), and tertiary stage (severe complications affecting organs) [4]. Other notable treponemal diseases include yaws. Yaws caused by *Treponema* subsp *pertenue* (TPE) is characterized by initial lesions appearing as painless ulcers on the skin, which can progress to more severe manifestations if untreated, leading to bone and cartilage deformities [5,6]. Bejel is another form, also known as endemic syphilis, caused by *Treponema* subsp. *Endemicum*, similar to yaws but primarily affecting mucous membranes and bones rather than the skin. Lastly, Pinta, a skin disease caused by *Treponema carateum* characterized chronic infections that can lead to severe complications if left untreated. Of all the Treponema species, yaw constitutes the highest burden of disease globally [7].

Non-human primates (NHPs) and humans are found to be affected with similar strains of *Treponema pallidum* subspecies *pertenue* (TPE), the strain that causes yaws in humans, with the potential for cross-transmission among them [8]. Studies shows, populations of wild Gorillas (*Gorilla gorilla gorilla*) and Chimpanzees (*Pan troglodytes troglodytes*) in the Democratic Republic of Congo, Gabon, Cameroon, and Guinea have been diagnosed with “Yaws” [9] characterized by yaws-like facial lesions resembling those observed in humans [10,11].

Infections in NHPs often manifest skin lesions similar to those in humans, including anogenital ulcers and orofacial lesions. The presence of *T. pallidum* in NHP populations raises concerns about potential zoonotic transmission to humans, especially in areas where human cases of yaws have been reported. The geographic distribution of *T. pallidum* infection among NHPs closely mirrors that of human yaws, suggesting a complex interplay between wildlife reservoirs and human health. In Tanzania, four (4) different NHP species, i.e., Olive baboons (*Papio anubis*), Yellow baboons (*Papio cynocephalus*), Vervet Monkeys (*Chlorocebus pygerythrus*), and Blue Monkeys (*Cercopithecus mitis*) were found to be infected with Treponema *TPE*, with widespread geographical distribution within the country [12]. The populations of wild baboons (*Papio anubis*) at Gombe National Park in Tanzania are naturally infected with *Treponema pallidum* subspp *pertenue* (TPE). However, the epidemiology and pathogenesis of the disease are not well known and hence the current study aimed at a better understanding of drivers and risk factors for recurrence and epidemiology of TPE infections in NHPs.

## Materials and Methods

### Study Area

The study was conducted at Gombe National Park located in northwestern Tanzania (4^0^ 41’S, 29^0^ 37’E), 16 km north of Kigoma town on the shores of Lake Tanganyika at an altitude of 765m above sea level. The National Park covers an area of 14 square miles (35.69 sq km) of land, and (since 2015) 20.72 km^2^ of water has been added from Lake Tanganyika, making an area of 56 square kilometers. The park comprises grasslands, woodlands, steep valleys, and tropical rainforest [13–15]

### Study design & Sample size

This study employed longitudinal design, using archived data (Health Monitoring records) from January 2019 to December 2024 to investigate disease risks, trends, and recurrence patterns. Additional data from March to June 2025 were incorporated into the dataset to provide a more comprehensive view of the current situation. We analyzed data collected from wild baboons (*Papio anubis*) during this period to examine the occurrence, dynamics, and trends of the disease. All archived records from 2019 to 2024 were analyzed to assess all potential disease risks, deaths and disappearances, injuries, interactions both within and between baboon troops as well as other animal species, Male intergroup immigrations and Effects of Treatments since some baboon troops have undergone antibiotic treatments at different times, administered both selectively to only affected individuals and more at some point broadly at the Troop level, in efforts to minimize the impact of recurring infections [16].

### Data collection

All data archived (January 2019 to December 2024), which includes monthly reports, necropsy reports, and baboons attendances on studied troops at Gombe Stream Research Centre, were used to study factors influencing infection rates in baboon troops. The recorded baboon data comprised of births, deaths, immigrations, illnesses and conditions, injuries, monthly diet, interactions between baboon troops and with other species, ranging behavior, and any encounter of unusual events where Presence of *Treponema* like clinical signs (ulcerative lesions, genital sores, or skin nodules) was recorded during routine behavioral follows.

### Description of Treponema clinical cases

In baboons, Treponema manifests as genital ulcerative or crusted lesions on the penis, scrotum, vulva, or perineum. Chronic lesions may be granulomatous or necrotic in appearance. Lesions may persist for months and may lead to scarring. Mucocutaneous Lesions: Less commonly, baboons can have lesions on the face, limbs, or oral mucosa. [17,18] At Gombe, affected individuals usually present the ulcerations on the anogenital area in females, for males ulcerations around the preputial opening that lead to phimosis and for females completely closure of the external vagina (vulva) and death may occur for untreated cases., mucocutaneous lesions have recently been observed, where facial (nose) and limbs were involved (Figure 1).

**Figure 1.**
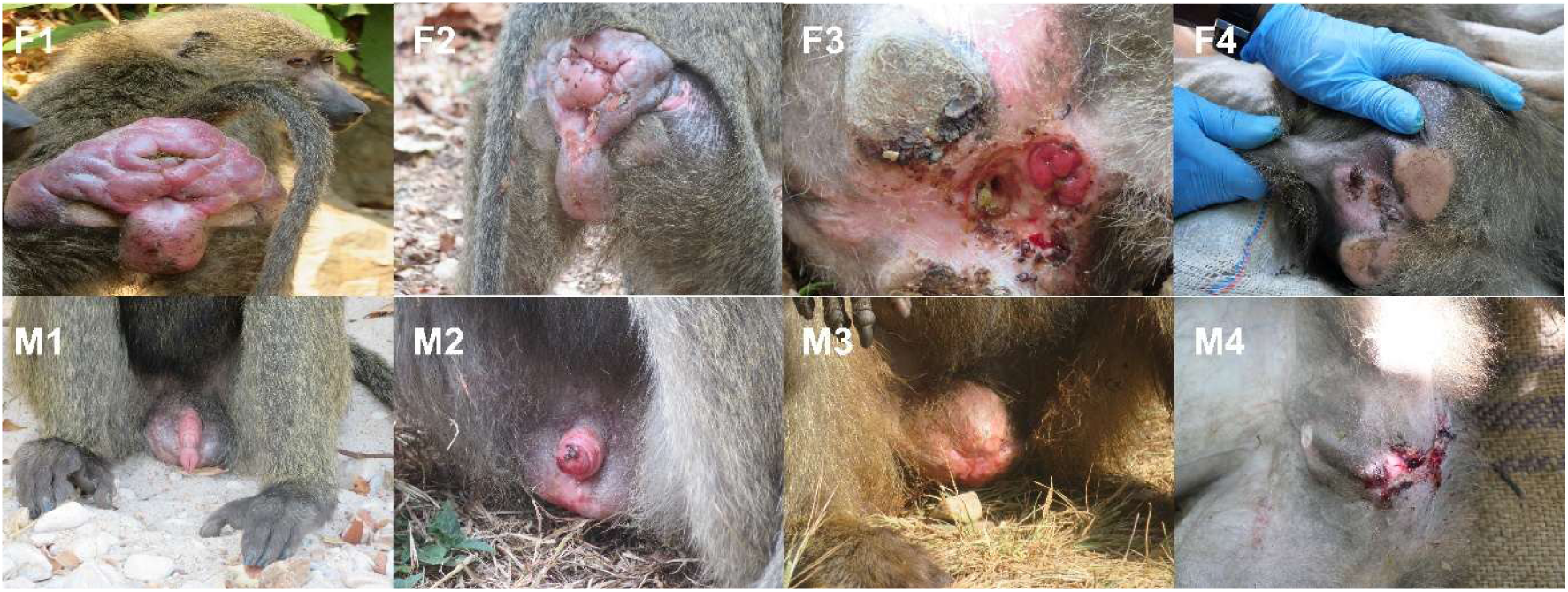
Baboons with different stages of Treponema Lesions - Left to Right, initial -> Severe cases as observed at Gombe National Park (Upper row Females (F1-F4), Lower Males (M1-M4).

### 3.0 Ethical clearance and research permits

This research was conducted in accordance with the Tanzania Wildlife Research Institute (TAWIRI) Guidelines for Conducting Wildlife Research (2012); http://tawiri.or.tz/wpcontent/uploads/2017/05/Wildlife-research-guideline.pdf). The approval was granted by TAWIRI and was approved by the Commission for Science and Technology (COSTECH), under Research Permit No. CST00001000-2024-2025-00245. Additionally, the study also adhered to the ethical guidelines outlined in the Code of Ethics of Sokoine University of Agriculture.

### Data analysis

Data was analyzed using R (version 4.5.1; R Core Team, 2025). Descriptive statistics were used to summarize demographic and epidemiological characteristics. Categorical variables (Sex, Age Category, Troop, Immigration status) were reported as counts and percentages. Disease prevalence (presence of Treponema signs) was calculated as the proportion of affected individuals and stratified by risk factors, including Sex, Age Category, Pregnancy state, Troop, Troop treatment, Immigration status, and Year. Statistical differences between troops were evaluated using chi-squared tests and summarized using the *gtsummary* package in R. Temporal trends in prevalence were visualized using line plots stratified by risk factors, generated with the ggplot2 package.

To identify independent risk factors, a Generalized Linear Mixed Effects Model (GLMM) with a logit link and binomial distribution was fitted to a longitudinal dataset comprising 17,946 monthly observations of approximately 390 baboons tracked from 2019 to 2024. Random intercepts for individual identity (’Baboon Name’) were included to account for repeated measures [19]. The model revealed substantial between individual variance (τ₀₀ = 8.92) and a residual variance of σ² = 3.29, yielding an intraclass correlation coefficient (ICC) of 0.73. Fixed effects accounted for 32.8% of the variance (marginal R²), while the full model including random effects accounted for 81.9% (conditional R²), indicating a strong influence of individual differences alongside significant fixed predictors (*p* < 0.001). Predictor variables included Sex, Age Category, Troop, Immigration status, Troop treatment, Presence of wounds, Pregnancy state, and Year. Results are reported as adjusted odds ratios (ORs) with 95% confidence intervals (CIs) and p-values using *gtsummary*. All categorical variables were treated as factors. The outcome variable was recorded as binary (1 = present, 0 = absent).

All analyses were conducted using complete case data; only observations with no missing values for the variables of interest were included. No imputation methods were applied to estimate or replace missing data.

The groups of Age Categories included were as follows: Adults comprised the group of baboons above 6 years old, Subadults (3-6 years old). Juveniles (Between 1 and 3 years old) and infants (All Below 1 year old) [19–21]. Seasonal analysis was based on two seasons of the year at Gombe, where the dry season being characterized by minimal precipitations below 100mm, food scarcity and high stress to the animals, the period spans from May to October. Wet season occurs from November to April, and is characterized by heavy rains 1,450 to 1,710 mm annually, which is the period where food is plentiful and conducive weather for the animals, a less stressful period [23,24]

## Results

### Baboon Population Structure and Dynamics

The study population consisted fluctuation number of around 230 habituated olive baboons (*Papio anubis*) over the study years (2019-2024), the populations had minor fluctuations in size based on births and immigration as observed during the study period (Figure 2).

**Figure 2.**
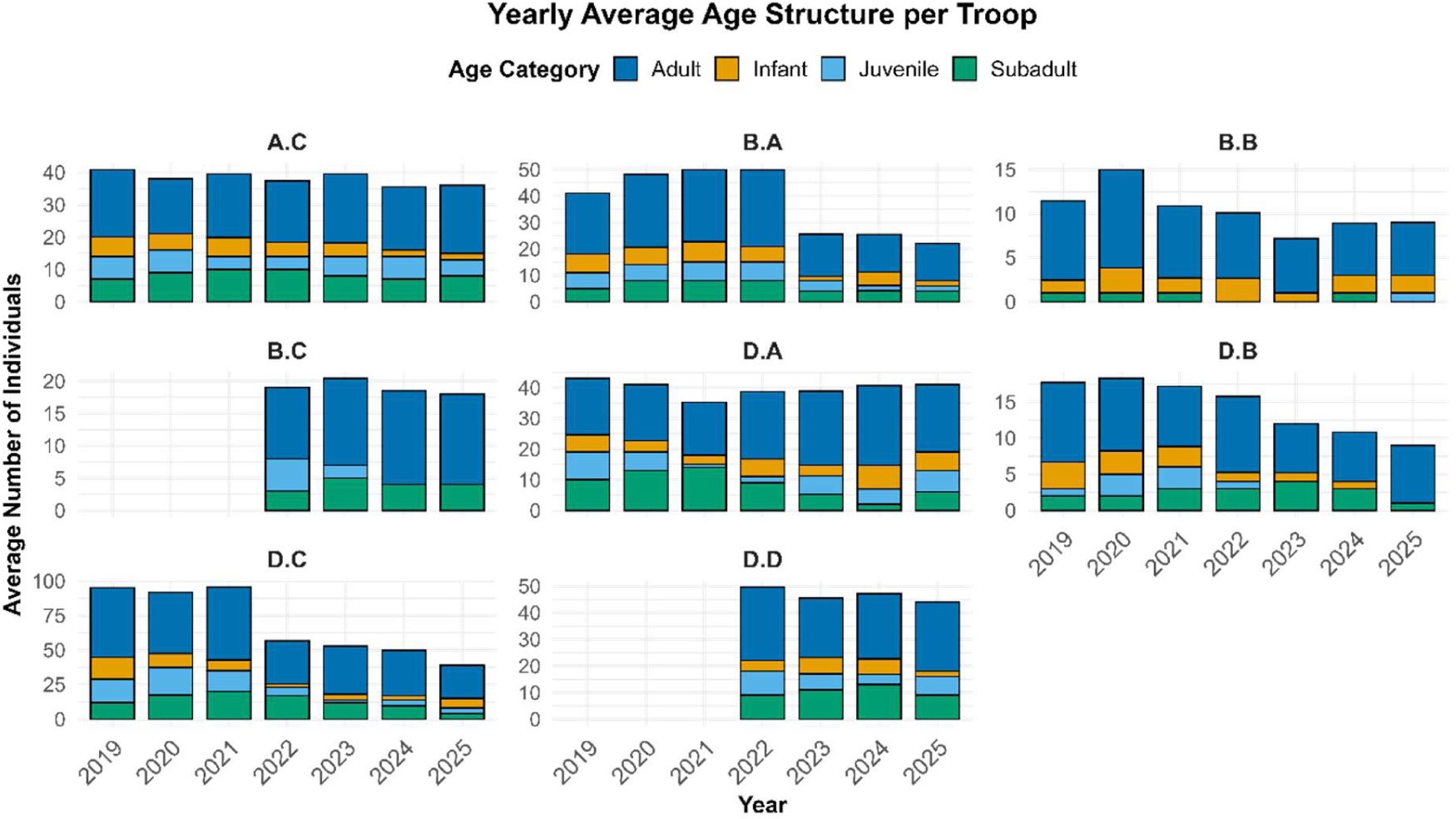
Population of Baboons study troops at Gombe National Park by age categories.

Over the study period, overall individual observations recorded monthly across all baboon troops were 17,946. These counts reflect repeated monthly observations of individuals rather than unique animals. The dataset was dominated by adult baboons, who accounted for 10,048 observations (56%), followed by subadults with 3,531 (20%), juveniles with 2,273 (13%), and infants with 2,094 (12%), which were initially distributed among six (6) baboon troops (A.C, B.A, B.B, D.A, D.B, and D.C). However, at the end of the Year 2021 and 2022 period, two of the troops (D.C and B.A) underwent fission events, giving rise to two additional troops D.D (from D.C) and B.C (from B.A), thereby increasing the total number of troops to eight [25] (Figure 2).

### Health Challenges Observed Among Baboons During the Study Period

General health conditions observed and reported during the study period included a range of disease symptoms and injuries, with notable variation across the study troops. Coughing was one of the most frequently reported symptoms among a few troops, primarily affecting Troop B.A where the prevalence was 0.1% (2/2912), A.C Troop 8.1% (226/2805), B.B Troop with approximately 38% (288/764), and D.B Troop with the highest observed prevalence at 51% (570/1,123) among all studied troops. Treponema cases accounted for most reported health conditions across all study troops, with an average prevalence of 11.24% as compared to all cases reported on each troop, as seen under Diseases and injuries, as summarized in Figure 3.

**Figure 3.**
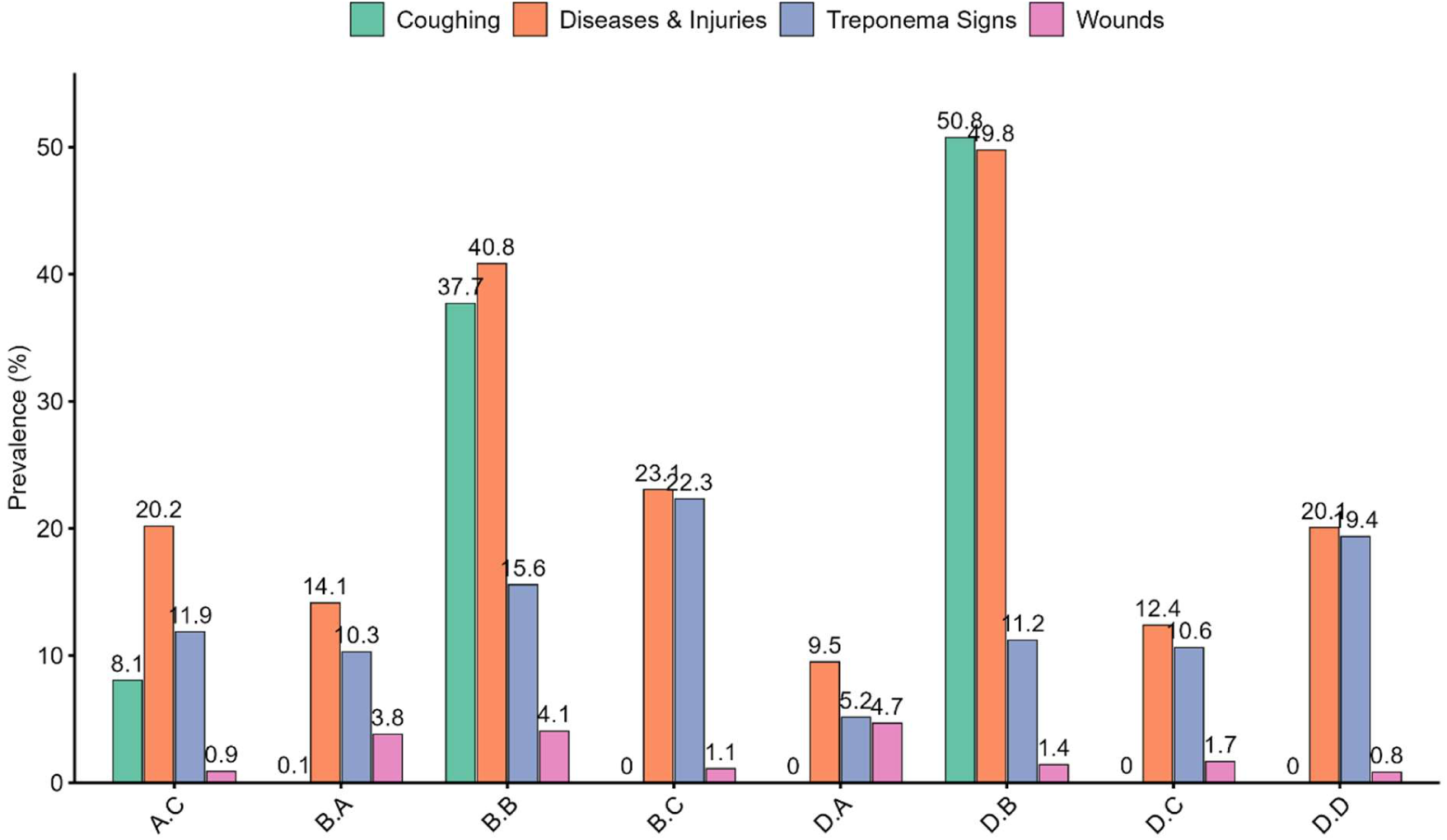
Health conditions reported to affect Baboons from January 2019 to December 2024 in the studied troops.

### Overall distribution of Treponema clinical Infection

Initial analyses using Chi-square tests suggested significant associations between Treponema clinical cases and named risk factors as listed on Table 1. However, given the repeated measures over the years and potential for non-independence among observations, we reanalyzed the data using Generalized Linear Mixed Models (GLMMs) to account for random effects. Most results were consistent with the Chi-square findings, indicating robustness of the observed trends.

**Table 1.** Overall distribution of clinical Treponema (Present vs Absent) across demographic and behavioral risk factors from January 2019 to December 2024.

| Risk Factor | 0<br>N = 15,928<sup>1</sup> | 1<br>N = 2,018<sup>1</sup> | p-value<sup>2</sup> |
| --- | --- | --- | --- |
| Sex |  |  | <0.001 |
| Female | 8,659 / 15,928 (54%) | 1,264 / 2,018 (63%) |  |
| Male | 7,088 / 15,928 (45%) | 754 / 2,018 (37%) |  |
| Unidentified | 181 / 15,928 (1.1%) | 0 / 2,018 (0%) |  |
| Age Category |  |  | <0.001 |
| Adult | 8,368 / 15,928 (53%) | 1,680 / 2,018 (83%) |  |
| Infant | 2,080 / 15,928 (13%) | 14 / 2,018 (0.7%) |  |
| Juvenile | 2,208 / 15,928 (14%) | 65 / 2,018 (3.2%) |  |
| Subadult | 3,272 / 15,928 (21%) | 259 / 2,018 (13%) |  |
| Troop |  |  | <0.001 |
| A.C | 2,472 / 15,928 (16%) | 333 / 2,018 (17%) |  |
| B.A | 2,612 / 15,928 (16%) | 300 / 2,018 (15%) |  |
| B.B | 645 / 15,928 (4.0%) | 119 / 2,018 (5.9%) |  |
| B.C | 421 / 15,928 (2.6%) | 121 / 2,018 (6.0%) |  |
| D.A | 2,756 / 15,928 (17%) | 150 / 2,018 (7.4%) |  |
| D.B | 997 / 15,928 (6.3%) | 126 / 2,018 (6.2%) |  |
| D.C | 4,772 / 15,928 (30%) | 568 / 2,018 (28%) |  |
| D.D | 1,253 / 15,928 (7.9%) | 301 / 2,018 (15%) |  |
| Seasons |  |  | 0.001 |
| Dry | 8,179 / 15,928 (51%) | 1,113 / 2,018 (55%) |  |
| Wet | 7,749 / 15,928 (49%) | 905 / 2,018 (45%) |  |
| Troop Treatment | 10,128 / 15,928 (64%) | 1,020 / 2,018 (51%) | <0.001 |
| Pregnancy | 457 / 15,928 (2.9%) | 22 / 2,018 (1.1%) | <0.001 |
| Wounds | 398 / 15,928 (2.5%) | 29 / 2,018 (1.4%) | 0.004 |
| Immigration | 565 / 15,928 (3.5%) | 68 / 2,018 (3.4%) | 0.7 |
| Interactions General | 775 / 15,928 (4.9%) | 174 / 2,018 (8.6%) | <0.001 |

<sup>1</sup>n / N (%)
<sup>2</sup>Pearson's Chi-squared test

During the study period, the overall prevalence of Treponema clinical signs consistent with Treponema infection across all study troops was 11.24% (2018/17946). Adults were the most affected age group, comprising 83% of all the cases, followed by subadults (13%), juveniles (3.2%), and infants (0.7%). Females exhibited a significantly higher prevalence than males (63% vs. 37%) Table 1.

Seasonal differences were observed, with more cases recorded during the dry season (55%) than the wet season (45%). Treatment was associated with a significant reduction in clinical signs, with treated troops showing lower prevalence compared to untreated ones (p < 0.001). Pregnancy was significantly protective against disease expression (p < 0.001), while the presence of wounds was positively associated with clinical signs (p = 0.003). No significant associations were found between infection prevalence and immigration events, mortality, or disappearances. The highest annual prevalence was recorded in 2022, accounting for 28% of total cases over the study period (p < 0.001) Table 1.

### Treponema distribution across Risk Factors

The mixed-effects logistic regression model (GLMM) was used to assess predictors of the presence of *Treponema* signs in baboons, accounting for repeated measures within individuals (random effect: “Baboon_Name”) [19]. Results indicated a significant variation existed between various demographics, temporal, and environmental risks factors as seen in Table 2, that were significantly associated with the risk of *Treponema* infection.

**Table 2.**
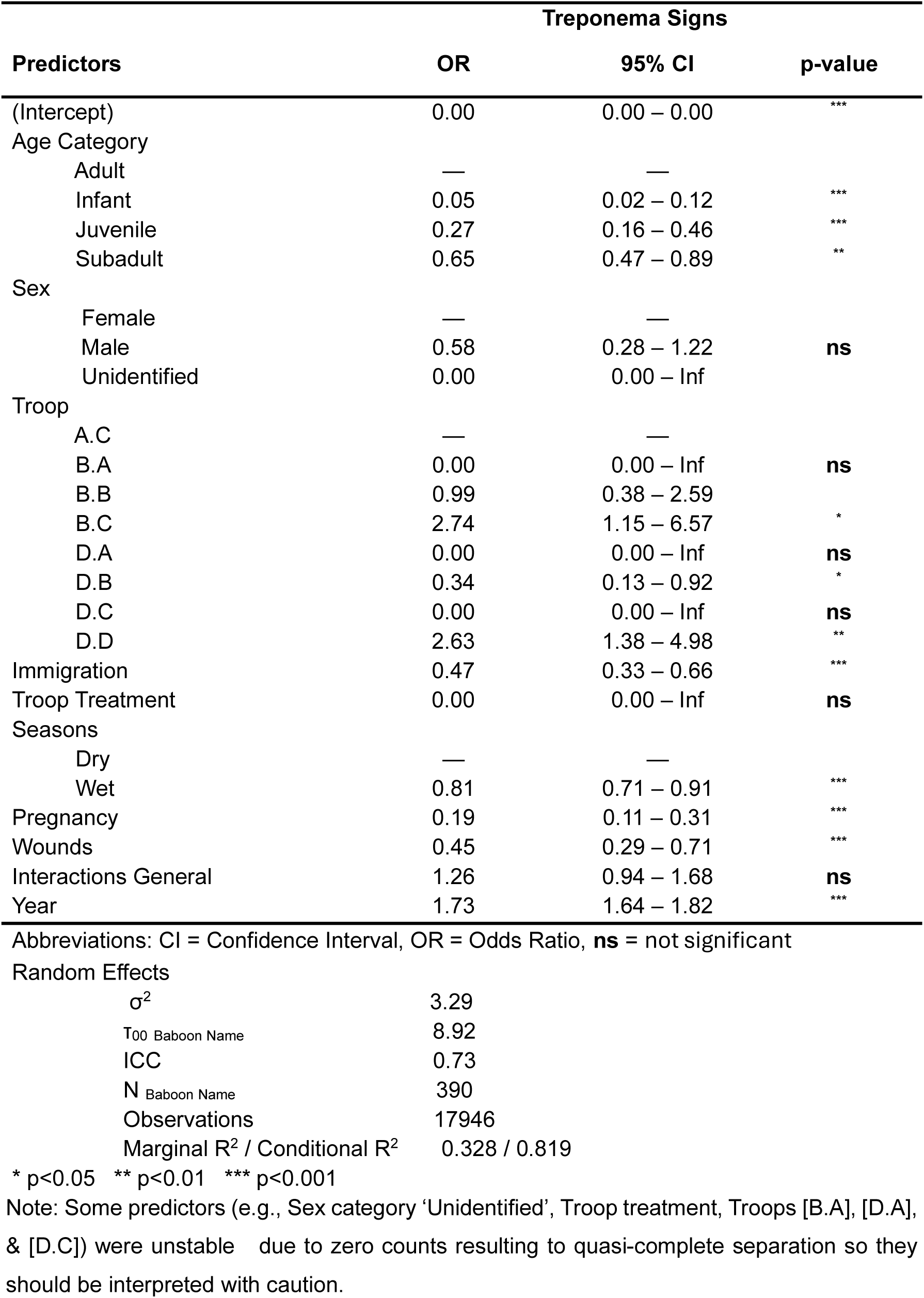
Predictors of Treponema Clinical Infection Signs in Wild Baboons Based on a Generalized Linear Mixed-Effects Model (GLMM) in Studied Baboon Troops at Gombe.

| Treponema Signs |  |  |  |
| --- | --- | --- | --- |
| Predictors | OR | 95% CI | p-value |
| (Intercept) | 0.00 | 0.00 – 0.00 | *** |
| Age Category |  |  |  |
| Adult | — | — |  |
| Infant | 0.05 | 0.02 – 0.12 | *** |
| Juvenile | 0.27 | 0.16 – 0.46 | *** |
| Subadult | 0.65 | 0.47 – 0.89 | ** |
| Sex |  |  |  |
| Female | — | — |  |
| Male | 0.58 | 0.28 – 1.22 | ns |
| Unidentified | 0.00 | 0.00 – Inf |  |
| Troop |  |  |  |
| A.C | — | — |  |
| B.A | 0.00 | 0.00 – Inf | ns |
| B.B | 0.99 | 0.38 – 2.59 |  |
| B.C | 2.74 | 1.15 – 6.57 | * |
| D.A | 0.00 | 0.00 – Inf | ns |
| D.B | 0.34 | 0.13 – 0.92 | * |
| D.C | 0.00 | 0.00 – Inf | ns |
| D.D | 2.63 | 1.38 – 4.98 | ** |
| Immigration | 0.47 | 0.33 – 0.66 | *** |
| Troop Treatment | 0.00 | 0.00 – Inf | ns |
| Seasons |  |  |  |
| Dry | — | — |  |
| Wet | 0.81 | 0.71 – 0.91 | *** |
| Pregnancy | 0.19 | 0.11 – 0.31 | *** |
| Wounds | 0.45 | 0.29 – 0.71 | *** |
| Interactions General | 1.26 | 0.94 – 1.68 | ns |
| Year | 1.73 | 1.64 – 1.82 | *** |

Abbreviations: CI = Confidence Interval, OR = Odds Ratio, **ns** = not significant

Random Effects
|  |  |
| --- | --- |
| $\sigma^2$ | 3.29 |
| T00 Baboon Name | 8.92 |
| ICC | 0.73 |
| N Baboon Name | 390 |
| Observations | 17946 |
| Marginal R<sup>2</sup> / Conditional R<sup>2</sup> | 0.328 / 0.819 |

\* p<0.05 \*\* p<0.01 \*\*\* p<0.001
Note: Some predictors (e.g., Sex category 'Unidentified', Troop treatment, Troops [B.A], [D.A], & [D.C]) were unstable due to zero counts resulting to quasi-complete separation so they should be interpreted with caution.

Troop B.C had 2.74 times higher odds of Treponema signs (p < 0.05), and Troop D.D had similarly elevated odds (OR = 2.63, p < 0.01) relative to the reference troop. Equally, Troop D.B had significantly lower odds (OR = 0.34, p < 0.05). Other troops showed zero or near-zero odds, likely due to zero counts in some categories, resulting in unstable estimates.

The Age category had a significant effect on distribution of *Treponema* clinical cases, Infants were much less likely to show signs (OR = 0.05, p < 0.001), followed by Juveniles (OR = 0.27, p < 0.001), and Subadults (OR = 0.65, p < 0.01) compared to adults. This indicates increasing risk with age i.e., Adults >> Subadults >> Juveniles >> Infants. Comparable trends were also observed using Chi-square test where adults accounted for most cases across all troops, with a statistically significant difference (p < 0.001) Table 1.

Sex: Male baboons showed lower odds of developing *Treponema* signs compared to females (OR = 0.58), indicating a 42% reduction in odds, but this effect was not statistically significant under GLMM model (CI: 0.28 –1.22) Table 2.

Season: Seasonal variation influenced infection prevalence, where the wet season was associated with a significant reduction in odds of *Treponema* signs (OR = 0.81, 95% CI: 0.71– 0.91, p < 0.001), relative to the dry season despite the wet season being longer (7 months) as compared to dry season (5 Months).

Pregnancy: Pregnancy was observed to be strongly protective, with pregnant individuals demonstrating an 81% reduction in odds of *Treponema* infection compared to non-pregnant counterparts (OR = 0.19, p < 0.001), indicating a potential protective or physiological effect during pregnancy.

Wounds: Baboons that were recorded to be wounded were significantly associated with reduced odds of Treponema signs (OR = 0.45, p < 0.001), which may reflect complex relationships between injury and infection risks.

Immigration status: Immigrant male baboons showed significantly lower odds of *Treponema* signs (OR = 0.47, p < 0.001), suggesting differences in exposure or susceptibility.

Troop Treatments: There was no significant difference between Treated and non-Treated Troops reflecting instability in the model likely due to sparse data or zero event counts in certain groups, and thus should be interpreted with caution

Year: A strong positive temporal trend was observed (OR = 1.73, p < 0.001), that marked an increase in infection risk over time with each subsequent year linked to a significant increase in odds of infection Table 2. indicating an overall upward trajectory in disease prevalence during the study period.

Random Effects and Model Fit: Significant individual-level variability was observed (τ₀₀ = 8.92), with an intraclass correlation coefficient (ICC) of 0.73, indicating that 73% of variation in Treponema signs was explained by differences between baboons. The model explained 32.8% of variance by fixed effects alone, increasing to 81.9% when including random effects.

### Trends of *Treponema* Infection across Risk Factors

#### Age Category

Throughout the study, a clear and consistent trend in disease prevalence emerged across age groups. Adults consistently showed the highest levels of prevalence each year, followed in order by Subadults, then Juveniles, and finally Infants Figure 4. This trend remained largely stable over time, indicating that older individuals were disproportionately affected. Notably, in Year 2022, the prevalence among Juveniles started to decline, with the prevalence in Year 2023 reaching levels nearly indistinguishable from those of Infants. The prevalence among infants remained consistently low, approaching zero throughout the entire study period. A brief period of overlap in prevalence between Subadults and Juveniles was observed in Year 2024, although Subadults generally maintained higher infection rates than younger individuals. Overall, Infants consistently showed minimal to no detectable prevalence, suggesting limited exposure or susceptibility within this youngest age group.

**Figure 4.**
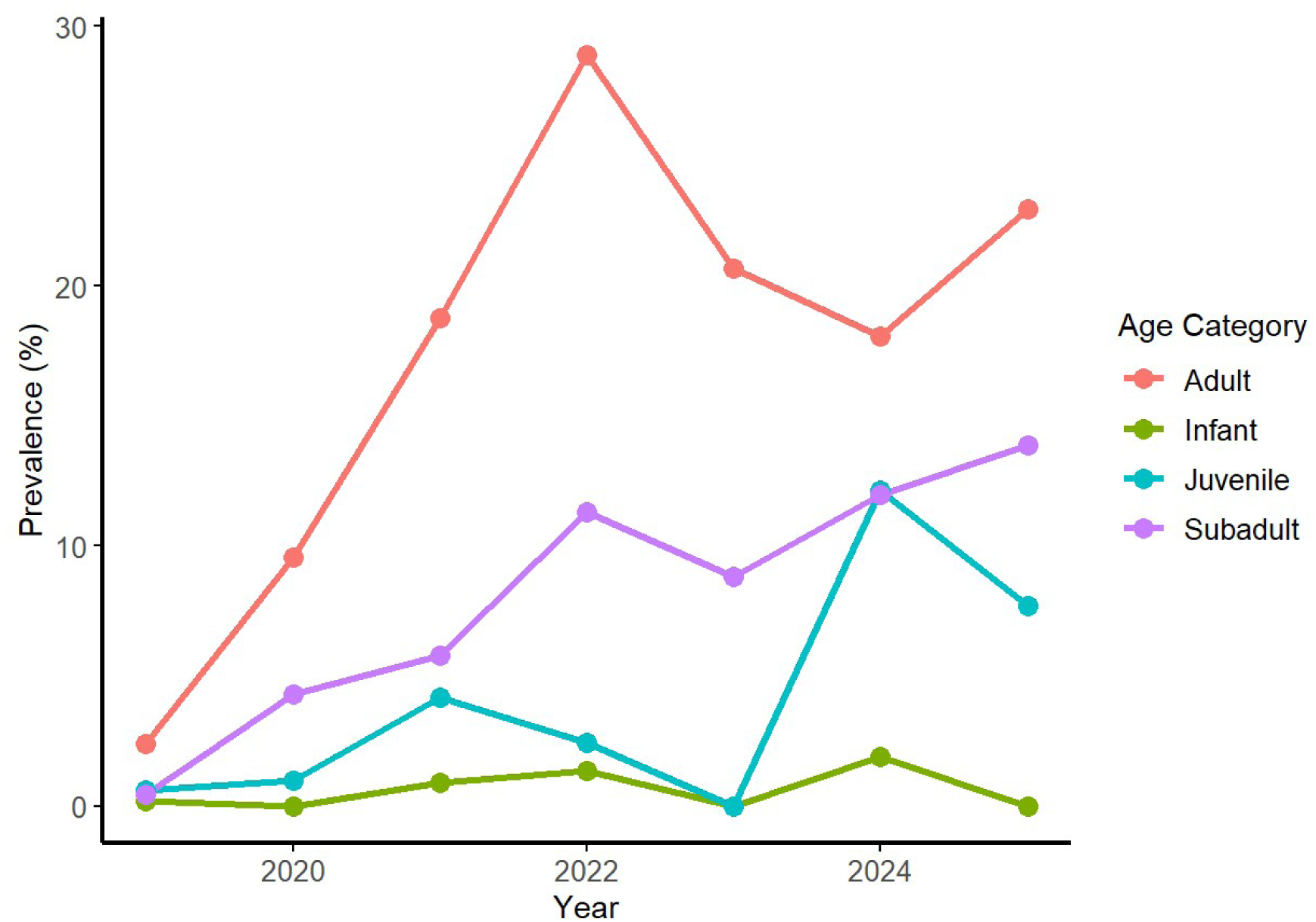
Prevalence of Treponema clinical infections across different age categories.

### Sex

Treponema clinical disease prevalence across sexes over the study period revealed a consistent trend of higher prevalence in females compared to males. From January 2019 through December 2024, females demonstrated a markedly higher risk for the infection, with annual prevalence rates exceeding those of males in each recorded year (Figure 5). This sex-based disparity was sustained with minimal year-to-year variability, indicating a stable and persistent pattern over time.

**Figure 5.**
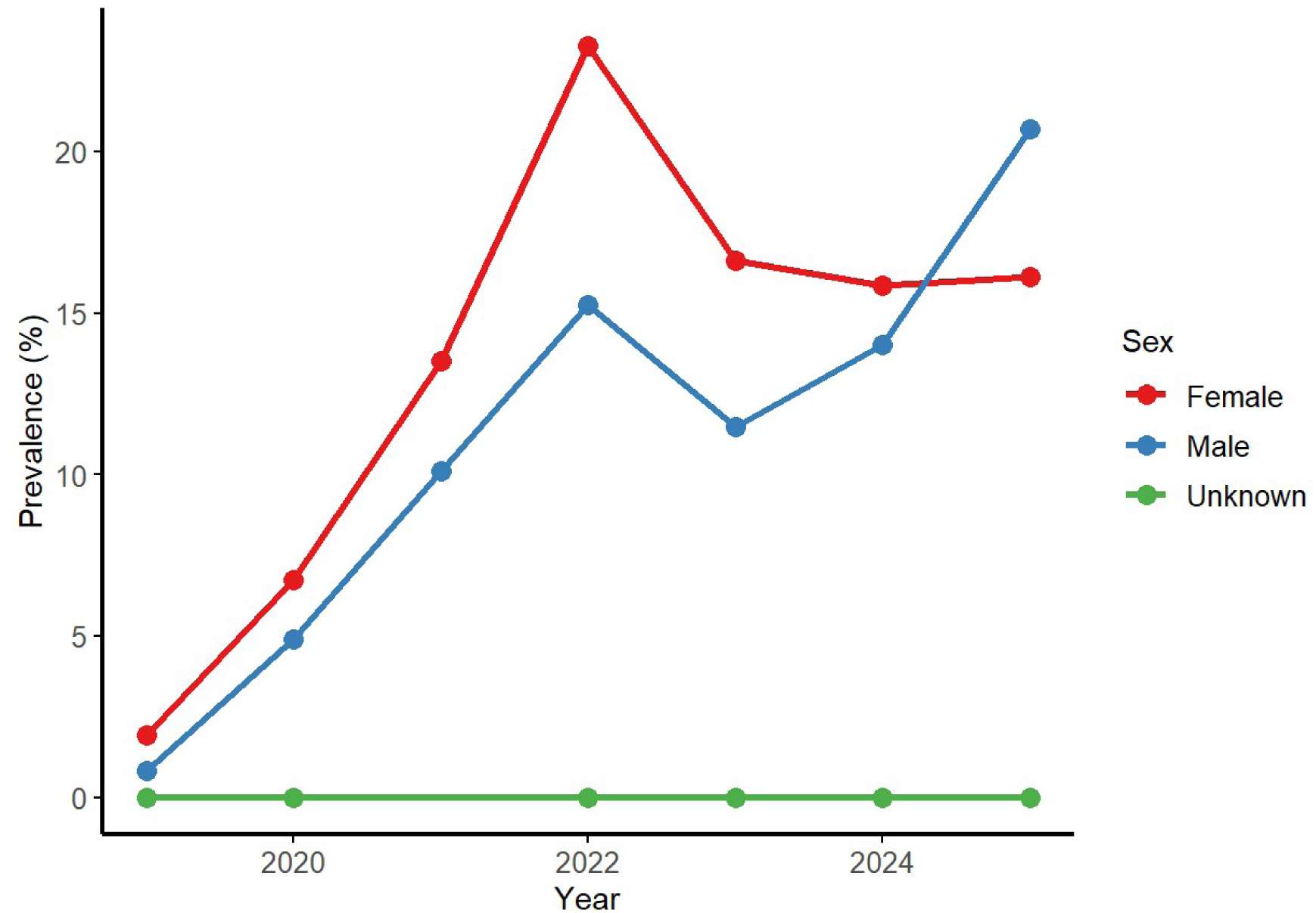
Trends of Treponema clinical infections by sex across Baboon troops.

However, an exception to this trend was observed in the most recent data snapshot from 2025. In this year alone, the prevalence in males slightly exceeded that in females, with the lines on the graph intersecting due to a marginal overlap in values. It is important to note that the 2025 data represent a preliminary cross-sectional snapshot rather than a fully aggregated annual dataset, and thus may not reflect the final trend for the current year.

Despite this single-year deviation, the overall trajectory supports a consistently higher prevalence of the disease in females.

### Troops

There was a consistent presence of Treponema-associated clinical infections across all studied troops throughout the study period. Troop DC exhibited a steady increase in clinical infection cases over the years, while troop DA maintained comparatively lower infection levels over time. Most troops experienced a notable surge in infection cases during 2022; however, troop BB deviated from this pattern, showing a continuous decline in infection rates beginning in 2021 and reaching the lowest levels by 2024.

The newly formed troops, DD and BC, exhibited the highest prevalence of Treponema infections by the end of the study period. Notably, at the time of their formation, both troops showed a modest reduction in infection cases relative to their respective parent troops, suggesting a temporary decrease in prevalence following troop division (Figure 6).

**Figure 6.**
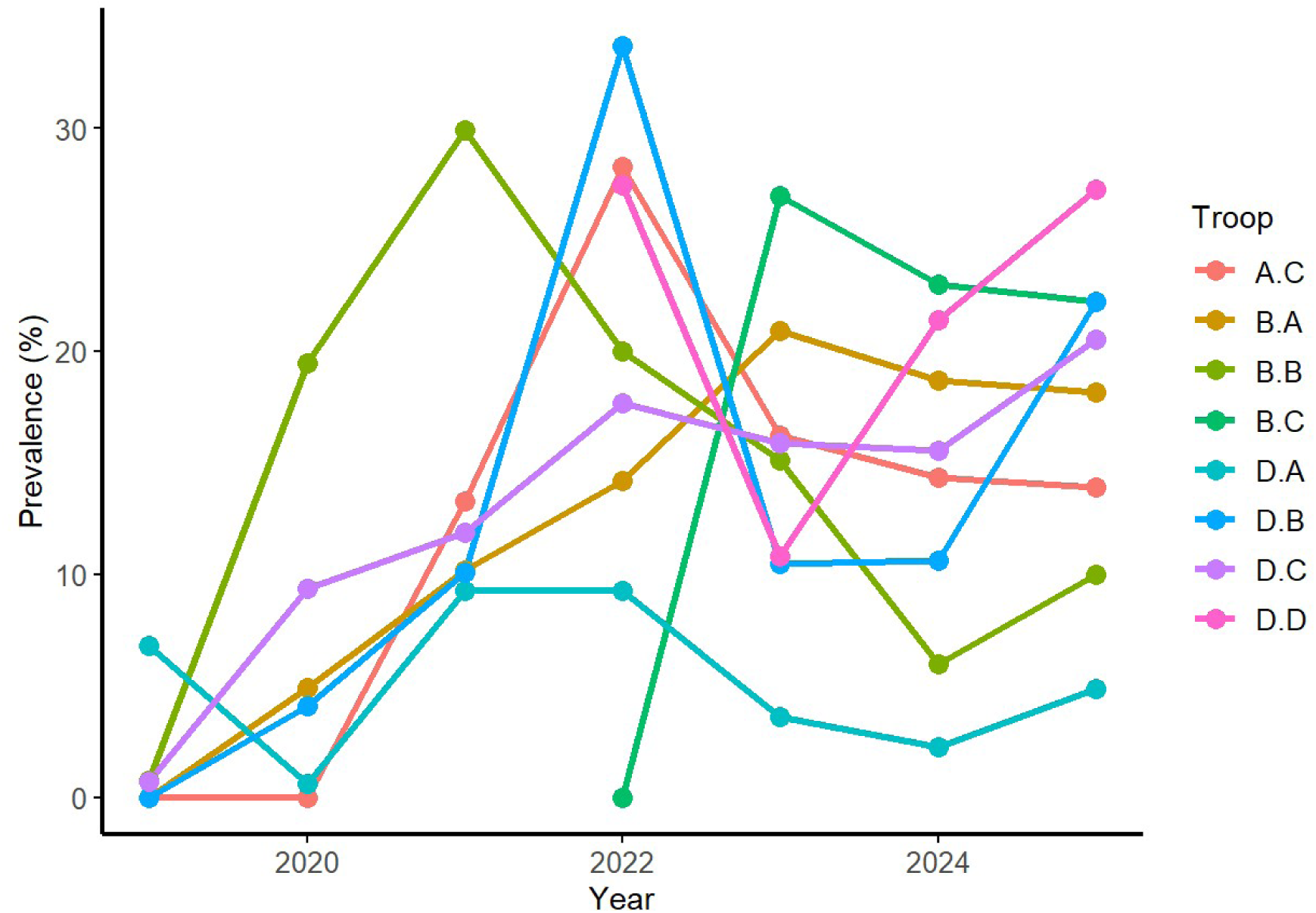
Trends of Treponema clinical infections across studied Baboons troops at Gombe.

### Seasonality

Analysis of the temporal distribution of clinical Treponema infections revealed a clear seasonal pattern, with most cases occurring during the dry season throughout the study period. This trend was consistent throughout all years, with the dry season exhibiting higher infection rates than the wet season. However, an exception to this pattern was observed in 2023, during which the wet season showed a slight but notable increase in infection cases, surpassing those recorded in the corresponding dry season. Despite this temporary deviation, the trend reverted in subsequent periods, with the dry season once again presenting higher infection rates than the wet season through to the end of the study period (Figure 7).

**Figure 7.**
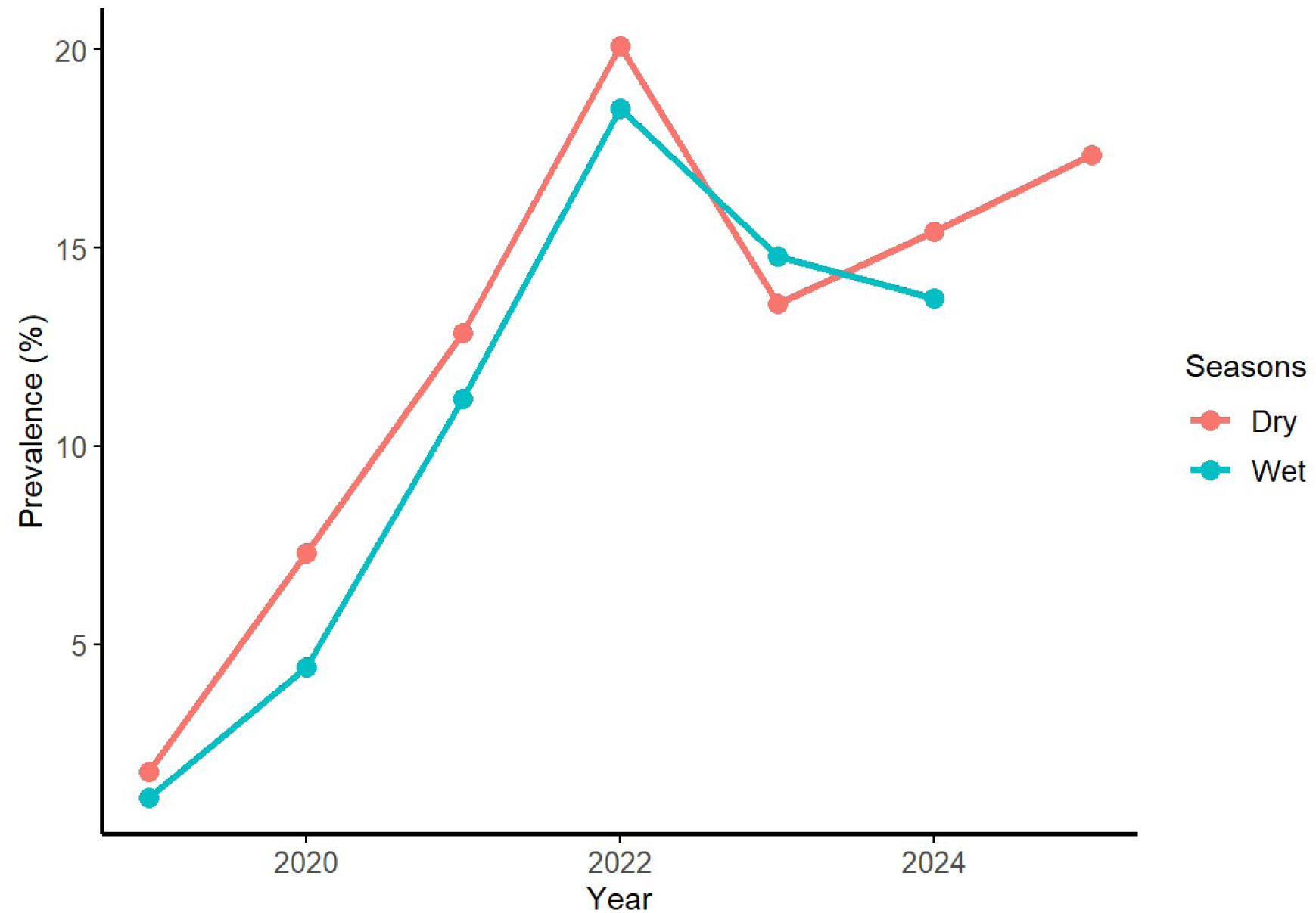
Trends of Treponema clinical infections over the years with different seasons.

## Discussions

At Gombe, Treponema infection has become endemic across all eight (8) baboon troops studied, spreading progressively over recent years despite extensive veterinary interventions aimed at eradicating the disease through treatment of the at-risk individuals in the affected troops and selective treatment of the affected individuals. The disease started to establish in the BA Troop in 2016 and expanded into neighboring troops, reaching all the troops through male migration as well as the ability of the pathogens to persist and latency on affected individuals who skipped medication during extensive treatments [2,25]. Since 1988, efforts have failed to achieve eradication, and recently, the disease has been associated with increased morbidity and mortality, particularly in older individuals. The dense social structure and frequent intergroup movement of affected males facilitate ongoing transmission, while repeated antibiotic use increases the risk of developing antimicrobial resistance.

### Infection rate in relation to risk factors

The results show a significant sex-based variation in *Treponema* infection among wild baboons, with females showing a markedly higher likelihood of infection than males. Even though females constituted 55% of the total sample, they accounted for 63% of infected individuals Table 1, where generalized linear mixed model (GLMM) also showed that male baboons had lower odds of developing *Treponema* signs (OR = 0.58), indicating a 42% reduction in odds, but this effect was not statistically significant (CI: 0.28 –1.22) Table 2. This disparity suggests that sex-specific behavioral or biological factors may influence transmission dynamics. Female baboons typically engage more frequently in grooming and maintain more stable group membership, as well as being constantly exposed to higher stress levels (Cortisol), potentially increasing cumulative exposure to pathogens [26–28]. Furthermore, sexual transmission of *Treponema pallidum* has been documented in non-human primates [29,30] and differences in genital morphology or mucosal vulnerability may contribute to increased susceptibility in females. Some studies reported a higher burden of genital lesions and active infections in adult females, supporting the patterns we observed in this study [31,32]. However, this finding differs from the observation on the Population of Gorillas in DRC, where males were associated with 1.9 times higher odds of showing Treponema signs (OR = 1.92, 95% CI: 1.15–3.22, *p* = 0.012) [11]. Hormonal and immunological differences between sexes, such as the influence of estrogen on immune function, may further modulate infection risk [33] These findings emphasize the importance of considering sex as a critical factor in the epidemiology of *Treponema* infections and call for further research into the behavioral and physiological mechanisms underlying this association.

The analysis also supports Age as a significant independent predictor of *Treponema* infection risks in wild baboons. The descriptive statistics showed a higher prevalence of infection among adults 83% and generalized linear mixed model (GLMM) also showed that the odds of infection increase markedly with age, independent of other variables. Non-adult age groups, infants, juveniles, and subadults had significantly lower odds of infection compared to adults. These findings suggest that transmission is likely driven by age-related behavioral and social factors, such as sexual activity or social interactions that are practiced with an increase in age. Several epidemiological studies of *Treponema pallidum* in non-human primates provide supportive evidence for our findings. A study done in Kenya showed that both adult and subadult olive baboons showed 100% seropositivity, while juveniles (∼61%) and infants (∼50%) showed progressively lower rates [29]. This gradient aligns with the GLMM results, indicating significantly lower odds of infection in younger age groups. Additionally, surveys across Tanzania have revealed infections among clinically asymptomatic individuals, suggesting latent or subclinical infections and underscoring the importance of adjusted analyses to reveal true age-related risk patterns [30]. Additional study across East Africa characterized more severe genital lesions in adults compared to subadults, which further reflects age-linked behavioral factors such as sexual activity as contributing to transmission [34]

We also found a significant seasonal pattern in *Treponema* infection among wild baboons, where the wet season was associated with reduction in probabilities of *Treponema* signs (OR = 0.81, 95% CI: 0.71–0.91, p < 0.001), relative to the dry season despite the longer period of wet season at Gombe (7 months) as compared to dry season. These findings suggest that environmental or behavioral factors associated with the dry season may facilitate transmission. Although targeted studies on the seasonality of treponemal infections in nonhuman primates are scarce. The observed pattern of elevated infection risk in the dry season may reflect factors such as increased stress during the dry season, which stimulates the emergence of the disease from latent state. We recommend future studies to examine seasonal and ecological drivers such as humidity, temperature, resource distribution, and social behavior to better understand the transmission ecology of *Treponema* in baboon populations.

Although male immigrants baboons comprised 3.5% of the overall sample, they accounted for a lower likelihoods of infection (3.4%) Table 1, as compared to non-immigrants baboons (OR = 0.47, p < 0.001) Table 2. The lower infection rate of male immigrants that suggest a protective association is a good to limit the spread of infections.

Pregnancy was significantly associated with a lower likelihood of *Treponema* infection in wild baboons. Although pregnant individuals comprised 2.9% of the total sample, they accounted for only 1.1% of infected cases (*p* < 0.001) Table 1. Under GLMM pregnant individuals demonstrated an 81% reduction in odds of *Treponema* infection compared to non-pregnant counterparts (OR = 0.19, p < 0.001), indicating a potential protective or physiological effect during pregnancy Table 2. This inverse relationship may reflect physiological or behavioral changes during pregnancy that reduce exposure risk or the development of clinical signs (latency). The finding suggests that pregnancy may offer temporary protection against the development of *Treponema* clinical infection, likely through reduced behavioral exposure or pregnancy-associated immune modulation. Future studies should explore infection dynamics across reproductive stages to better understand host susceptibility and transmission patterns.

Effect of Troop treatments, under descriptive statistics, we observed a notable difference where 62% of individuals belonged to treated troops, and within this group, 64% were uninfected, compared to 51% uninfected in untreated troops. This difference was statistically significant (*p* < 0.001), Table 1, suggesting a potential protective effect of treatment at the troop level. However, analysis using GLMM showed no significant difference due to zero event counts in certain Troops reflecting instability in the model and thus should be interpreted with caution Table 2. These findings suggest that troop-level treatment alone does not independently predict *Treponema* infection, once individual level variables are considered. The crude effect observed in descriptive statistics was likely driven by age or other demographic imbalances between treated and untreated troops.

There was a prominent upward trend in *Treponema* infection prevalence among baboons over the years from 2019 to 2024, with infected cases rising from 2.2% in 2019 to a peak of 28% in 2022, followed by a slight decline to 20% in both 2023 and early 2025 (though the 2025 dataset remains small) (*p* < 0.001). The GLMM model indicates a consistent annual increase in infection risk, A positive temporal trend (OR = 1.73, p < 0.001), indicating an overall upward trajectory in disease prevalence during the study period. Emphasizing a meaningful temporal escalation. This temporal acceleration may reflect cumulative transmission dynamics within stable social groups, with increasing pathogen circulation over time. Primates’ studies in Uganda and Rwanda showed that seropositive rates reached approximately 33% across 9 non-human primate species during surveillance of yaws-like outbreaks, where baboons had higher prevalence of 76.9% [35]. Likewise, long-term surveillance in East Africa has revealed chronic, widespread *Treponema pallidum* infections across multiple nonhuman primate taxa, suggesting enduring endemicity and potential expansion over time [29] Furthermore [30], *Treponema* infection to be widespread across the country. The study emphasizes the importance of monitoring trends, including serology and clinical assessment combined with Genetic strain tracking to unravel whether the observed increase reflects expansions in transmission, pathogen evolution, or changing host environment exchanges.

## Conclusion

Our study discloses that *Treponema* infection in wild baboons is influenced by multiple demographic, behavioral, and ecological factors. Age emerged as the strongest predictor, with adults showing significantly higher odds of infection, consistent with a cumulative exposure model. Females were more likely to be infected than males, likely reflecting sex-specific behaviors or physiological susceptibility, like exposure to higher levels of cortisol (stress). Seasonal variation showed a higher infection rate during the dry season, and Male immigrant individuals had significantly lower odds of infection, suggesting limited early exposure. Pregnant baboons are observed to be less likely to develop sign of infection, possibly due to the disease going into latency during the gestation period or reduced exposure risk. Lastly, infection prevalence increased markedly over the study years, indicating ongoing transmission and possible expansion. Together, these findings underline the complex interplay of host biology, behavior, and environment in shaping *Treponema* dynamics, and highlight the need for continued longitudinal monitoring and targeted studies on transmission pathways.

## Acknowledgement

The authors thank the Jane Goodall Institute-Tz, Tanzania Commission for Science and Technology (COSTECH), Tanzania Wildlife Research Institute (TAWIRI) and Tanzania National Parks (TANAPA) for support and providing permits for the long-term research at Gombe, we are also grateful to the work of all Gombe Stream Research Centre staff, for this work specifically baboon researchers without whom this work would not be possible.

## Author Contributions

Conceptualizing of research idea, writing of proposal, data collection, and drafting of manuscript DM. Supervision and assists in the field during data collection, AC, JR Supervision of the whole research, data analysis and interpretation, review and perfection of manuscript PW and AH. All the authors have read and agreed to the published version of the manuscript.

## Data availability

All data are included in this article.

